# Inflammasome mediated neuronal-microglial crosstalk: a therapeutic substrate for the familial *C9orf72* variant of the frontotemporal dementia/amyotrophic lateral sclerosis

**DOI:** 10.1101/2022.01.27.478078

**Authors:** Kyle J. Trageser, Chad Smith, Eun-Jeong Yang, Ruth Iban-Arias, Tatsunori Oguchi, Maria Sebastian-Valverde, Umar Haris Iqbal, Henry Wu, Molly Estill, Md Al Rahim, Urdhva Raval, Francis J Herman, Yong Jie Zhang, Leonard Petrucelli, Giulio Maria Pasinetti

## Abstract

Intronic G_4_C_2_ hexanucleotide repeat expansions (HRE) of *C9orf72* are the most common cause of familial variants of frontotemporal dementia/amyotrophic lateral sclerosis (FTD/ALS). G_4_C_2_ HREs in *C9orf72* undergo non-canonical repeat-associated translation, producing dipeptide repeat (DPR) proteins, with various deleterious impacts on cellular homeostasis. While five different DPRs are produced, poly(glycine-arginine) (GR) is amongst the most toxic and is the only DPR to accumulate in the associated clinically relevant anatomical locations of the brain. Previous work has demonstrated the profound effects of a poly(GR) model of *C9orf72* FTD/ALS, including motor impairment, memory deficits, neurodegeneration, and neuroinflammation. Neuroinflammation is hypothesized to be a driving factor in the disease course; microglia activation is present prior to symptom onset and persists throughout the disease. Here, using an established mouse model of *C9orf72* FTD/ALS we investigate the contributions of the nod-like receptor pyrin-containing 3 (NLRP3) inflammasome in the pathogenesis of FTD/ALS. We find that inflammasome-mediated neuroinflammation is increased with microglial activation, cleavage of caspase-1, production of IL-1β and upregulation of *Cxcl10* in the brain of *C9orf72* FTD/ALS mice. Excitingly, we find that genetic ablation of *Nlrp3* significantly improved survival, protected behavioral deficits and prevented neurodegeneration suggesting a novel mechanism involving HRE-mediated induction of innate immunity. The findings provide experimental evidence of the integral role of HRE in inflammasome-mediated innate immunity in the *C9orf72* variant of FTD/ALS pathogenesis and suggest the NLRP3 inflammasome as a therapeutic target.

## Introduction

Amyotrophic lateral sclerosis (ALS) is a progressive neurodegenerative disorder in which patients exhibit selective loss and demyelination of corticospinal motor neurons leading to terminal motor function impairments [1]. The pathophysiology underlying ALS is poorly characterized, which leaves patients with limited therapeutic options. *C9orf72* G_4_C_2_ repeat hexanucleotide expansions (HRE) represent ALS’s most common genetic cause and frontotemporal dementia (FTD), which represents approximately 25 % of familial ALS cases [2]. Intronic G_4_C_2_ HRE of *C9orf72* are the most common feature of the *C9orf72* familial variants of FTD/ALS resulting in the generation of several neurotoxic dipeptide repeat (DPR) proteins [3]. Most importantly, the poly (glycine-arginine) (GR) is amongst the most toxic and the primary DPR to accumulate in susceptible brain regions undergoing degeneration in the brain of ALS/FTD subjects [4]. Recent evidence strongly supports the pathogenetic role of poly (GR) in *C9orf72* FTD/ALS and models of *C9orf72* FTD/ALS, including motor impairment, memory deficits, neurodegeneration, and neuroinflammation [5,6].

Here we find evidence suggesting a potential new mechanism by which neuronal stress is induced by DPR proteins in the *C9orf72* variant of FTD/ALS. Most important, we find that silencing *Nlrp3*-mediated innate immunity significantly attenuates mortality and neurodegeneration and most notably mitigates behavioral impairment. The findings provide evidence of the integral role of innate immunity in c9FTD/ALS pathogenesis and suggest the NLRP3 inflammasome as a therapeutic target.

The high rate of concurrent diagnosis of FTD and ALS and the ongoing investigation into overlapping pathomechanisms have led to the notion of an FTD/ALS neurodegenerative spectrum disorder [7]. Recent investigations into the pathogenesis of ALS find that dysregulated innate immune activity may act as a detrimental mechanism that affects neuronal structure and function in line with our findings [8]. Our evidence suggests that activation of innate immunity in our model system [9], is highly relevant for its deleterious effect in promoting neuroinflammatory cascades through a mechanism involving proinflammatory cascades in the brain, as also described in other neurodegenerative conditions [10–12]. For example, since neuronal stress is a common feature in other forms of FTD/ALS, our evidence provides the basis for the broad therapeutic translatability in other forms of ALS, including SOD1^G93A^ and TDP-43.

Collectively, our study adds critical information toward understanding the etiology and pathogenesis of the familial *C9orf72* variant of FTD/ALS implicating novel mechanisms involving chemokines-mediated activation of the NLR3P inflammasome mediated innate immunity as a therapeutic target for the *C9orf72* variant of FTD/ALS.

## Materials and Methods

### Reagents and mice

The AAV_9_-GFP and AAV_9_-GFP-(GR)_100_ viruses were a kind gift from Dr. Leonard Petrucelli. Unless specified otherwise, cell culture reagents were obtained from ThermoFisher Scientific (Waltham, MA) and antibodies were obtained from Abcam (Cambridge, MA). C57BL/6J WT mice (#000664) and *Nlrp3*^−/−^ mice (#021302) were obtained from The Jackson Laboratory (Bar Harbor, ME) and socially housed on a 12:12-h light/dark cycle with lights on at 07:00 h in a temperature-controlled (20□±□2□°C) vivarium. Mice were given food and water ad libitum and were bred to obtain mouse pups used in our studies. All procedures were approved by the Institutional Animal Care and Use Committee of the Icahn School of Medicine at Mount Sinai.

### *C9orf72* experimental model

Upon confirmation of pregnancy, WT and *Nlrp3*^−/−^ dames were separated and monitored daily for litters. After the identification of a milk spot to confirm that newborn (p0) pups were being nursed, the pups were injected with AAV_9_-GFP or AAV_9_-GFP-(GR)_100_ by the ICV method [13]. Pups were cryoanesthetized by placing them in a padded 15 mL tube partially submerged in a slurry of water and crushed ice. Upon confirmation of anesthesia by toe-pinch, we identified the point of injection, approximately two-fifths of the distance between bregma and each eye. A Hamilton syringe (#7634-01; Reno, NV) with a 32 gauge needle (10 mm long, Hamilton #7803-04) was loaded with 4 µL of the AAV vector and placed 2 mm deep, orthogonal to the mouse’s head at the point of injection. 2 µL of the AAV vector was injected at a rate of 1 µL min into each lateral ventricle [14]. Upon completion of injections, each pup was rapidly returned to physiological temperatures by placing on a warming pad. The litters were returned to dames upon completion of injection for the entire litter and monitored for acceptance. Mice were weaned at p30 by sex. For confirmation of diffusion prior to studies, a small number of mice were injected with Trypan Blue and sacrificed two hours later for dissection of the brain and examination by stereoscopic microscope.

### Behavioral testing

After reaching 3 months of age, mice were tested through a battery of behavioral tests. All behavioral testing took place during the light phase of the day. On all days of behavioral testing, mice were acclimated to an anteroom directly adjacent to the behavioral testing room for 30 min. On Day One, mice were tested in the Open Field Test for basal anxiety and general locomotor impairments [15]. The Open Field apparatus consisted of a 40cm x 40cm x 40cm Plexiglas box with opaque white walls, situated within a dimly-lit room (200 lux). Mice were placed in the center of the apparatus and were allowed to freely explore for 10 minutes before being returned to their home cage. On Day Three through Five, mice were tested in a Hanging Wire Test for muscular impairments [16]. The Hanging Wire apparatus consisted of a 2 mm-thick wire suspended 35 cm over a layer of corn cob bedding, situated in a brightly lit room (500 lux). Mice were lifted from their home cage by the base of their tail and were placed near the wire until they grasped it with their forelimbs. The number of falls over a 2 min period was recorded. Falls in which the mice hung from the wire from their hindlimbs were excluded from the number of falls. At the end of the behavioral trial, mice were returned to their home cages. On Days Seven through Nine, mice were tested in a Contextual and Cued Memory Test in two Contexts. Context A was a 30cm x 24cm x 21cm conditioning chamber (Med Associates, Fairfax VT) within a room with white walls and bright lighting. Context A had a bare metal grid floor, bare grey walls, bright lighting, and background fan noise. Context A was cleaned with a 0.5% hydrogen peroxide solution (Virox, Oakville Canada) between each trial. Context B was a chamber of the same dimensions within a room with dim lighting. The chamber had a white plastic floor, curved white plastic walls, dim lighting, no background fan noise, and was scented with 0.25 % benzaldehyde in 70 % ethanol. Context B was cleaned with a 70 % ethanol solution between each trial. On Day Seven, mice were allowed to explore Context A for 4 min. At 180s, white noise (85 dB) played for 30 s and was co-terminated with a footshock (2 s, 0.75 mA). After the 4 min trial, the mouse was returned to its home cage. On Day Eight, mice were allowed to explore Context A for 4 min in the absence of white noise and footshock (Context Recall). On Day Nine, mice were allowed to explore Context B in the constant presence of white noise (85 dB, Cue Recall). For all tasks, the behavior was analyzed and recorded with Any-Maze v6.0 (Stoelting, Wood Dale IL).

### Immunohistochemistry

After completion of behavioral studies, a subset of mice was deeply anesthetized with ketamine/xylazine (100 mg/kg + 10 mg/kg, intraperitoneally), and then perfused transcardially with cold sterile PBS followed by 4 % PFA in PBS. Brains were removed, drop-fixed in 4 % PFA overnight, then washed once with cold PBS and stored in PBS. Tissue sections (50 μm thick) were taken with a vibratome (Leica; Wetzlar, Germany) and were stored in PBS with 0.02 % sodium azide (w/v). Sections were washed in PBS followed by 10 min permeabilization in 0.1 % Triton X-100 in PBS (PBST). Sections were then incubated in blocking solution (5 % normal goat serum in PBST) for 1.5 hr. The sections were washed three times with PBST and incubated with primary antibodies diluted in blocking solution: 1:500 Rabbit anti-Iba1 (ab178846, Abcam) and 1:250 chicken anti-GFP (A10262, ThermoFisher) overnight at 4 °C. Post incubation, the sections were washed three times in PBST and incubated with secondary antibodies diluted in blocking solution for two hours at room temperature (RT): Goat anti-Rabbit conjugated Alexa Fluor 568 (Abcam #175471, 1:500) and 1:500 Goat anti-Chicken Alexa Fluor 488 (A11039, Thermofisher). The sections were washed three times in PBST and incubated in 1μM DAPI solution (#ab228549, Abcam) for 5 min. The sections were washed twice in PBS and mounted on slides using ProLong Diamond Antifade Mountant (P36970, ThermoFisher).

### Stereology and cell morphology

Assessment of neuronal loss was assessed by GFP-expressing neurons count in the in the layers IV and V of the primary somatosensory (S1) cortex. Similarly, microglia density was determined in the same layers IV and V of the S1 cortical region by MBF Stereo Investigator (Williston VT). The number of sections required for each region was determined by an initial pilot study which found that six sections were needed for the S1 cortical region, each spaced equidistantly for both GFP-expressing neurons and IBA-1 immunopositive microglia. A widefield microscope (AxioImager M2/Z2, Carl Zeiss, Oberkochen Germany) along with MBF Stereo Investigator were used to visualize each section and to determine the number of microglia in each region. Using the Optical Fractionator workflow, the region of interest was traced onto each section based on the mouse brain atlas by Paxinos and Franklin. This was followed by a systematically random grid overlay consisting of squares measuring 300 µm by 300 µm, covering the region of interest. From each of these squares, an unbiased sampling region (100 µm by 100 µm) was used to manually count microglia somata in the area. Once each section was completed, the software ran an algorithm to estimate the number of microglia in the region. In parallel for IBA-1 immunopositive microglia, the Cavalieri estimator in the software was used to determine the volume of each region. The microglia count and volume were then used to determine the microglia density in the S1 cortical region. For assessment of cell morphology, images of coronal sections were taken on a Zeiss LSM880 Airyscan confocal microscope (Oberkochen, Germany) using an X20/0.8 NA air immersion objective controlled by Zeiss Zen Black software. For 3D analysis, z-stack images were obtained by capturing an image every 0.7μm covering the entire 50μm-thick section. Images were deconvoluted using AutoQuant X3.1 (Media Cybernetics, Rockville MD) and 3D analysis was performed using Imaris 9.1.2 (Bit Plane Inc, Concord MA) using the surface tool to reconstruct the soma and the filaments tool to reconstruct the branches.

### Western blotting

After completion of behavioral trials, a subset of mice was sacrificed and brain regions were immediately frozen on dry ice. Cerebral cortex was lysed with 1X Cell Lysis Buffer supplemented with 1 mM PMSF (Cell Signaling #8553S, Danvers MA) and Protease Inhibitor Cocktail (Sigma-Aldrich #11873580001). Tissue lysates were separated by electrophoresis and then transferred to a nitrocellulose membrane. Membranes were blocked for 1 h at RT, followed by treatment with primary antibodies (anti-caspase-1 antibody, 20B-0042; anti-IL-1β antibody 6243S; anti-α-tubulin antibody, T9026) at 4°C for overnight. Each membrane was then treated with a secondary antibody conjugated with horseradish peroxidase (anti-rabbit-or anti-mouse IgG-HRP-antibodies) for 1h at RT. Each band was detected using chemiluminescence detection kit (32106; Thermo Fisher Scientific) and data was analyzed using image J software to measure relative protein expression.

### Cortical microglia cultures

Cortices from 1-3 day-old C57BL/6J mouse pups were isolated, digested, and seeded at a density of 8 cortices per 24-well plate (12ml total volume). Every three days, the medium (DMEM + 10 % FBS + 1 % penicillin-streptomycin) was replenished. After 3 weeks, mixed glial cultures reached confluence and were isolated by mild trypsinization as previously described [17]. Briefly, cells were washed with culture medium without FBS and treated with a mixture of trypsin (0.25 % without EDTA) and DMEM-F12 medium in a 1:3 ratio. After 40 min incubation, mixed glial cells detached and left a layer of microglia attached to the bottom of the culture dish. Pure microglia were isolated by 15 min incubation with trypsin (0.05% with EDTA) at 37°C followed by gentle shaking. Cells were counted and seeded in 24-well plates at a density of 7.5 × 10^4^ cells/well.

### Cortical neuron cultures

Cortices from 3-day-old C57BL/6J mouse pups were isolated and finely diced in ice-cold HBSS. Cortices were then incubated in 10X Trypsin Solution (Sigma #59427C) with DNase I (Sigma #D4513) for 15 min at 37 °C with period inversion. Cells were then spun at 200 g for 5 min, and the pellet was triturated with a serological pipette and strained in a 40 μm cell strainer (BD Falcon #352340). Centrifugation followed by trituration was repeated once, and then the pellet was spun down and resuspended in Neurobasal medium supplemented with 0.25 % GlutaMAX, 2 % B-27, 10 % Fetal Bovine Serum, and 1 % Penicillin/Streptomycin at a density of 5 × 10^5^ cell/mL. Two hours later, the medium was changed to Neurobasal medium supplemented with 0.25 % GlutaMAX, 2 % B-27, and 1 % Penicillin/Streptomycin, and half the medium was replenished every 3-4 days afterward. Three days later, cells were infected with 2 × 10^10^ vg/mL AAV. Expression was confirmed by fluorescence microscopy. Cell supernatant and lysates were collected for assessment of LDH release and CXCL10 production (R&D Systems #DY466).

### Analysis of Cytokines

IL-1β in microglia culture supernatant was measured with Mouse IL-1β /IL-1F2 DuoSet ELISA Kit (R&D Systems DY401, Minneapolis MN) according to the manufacturer’s instructions. CXCL10 was measured in neuronal culture supernatant with Mouse CXCL10 DuoSet ELISA (R&D Systems DY466-05, Minneapolis MN) according to the manufacturer’s instructions. Primary microglia were stimulated with recombinant Mouse CXCL10 protein (R&D Systems 466-CR-608 050/CF, Minneapolis MN) and incubated with Mouse CXCL10 antibody (R&D Systems AF-466-609 NA, Minneapolis MN). TNF-α and IL-1β were measured in the supernatant by ELISA with Mouse TNF-α Quantikine ELISA Kit (R&D Systems MTA00B, Minneapolis MN) and Mouse IL-1β/IL-1F2 DuoSet ELISA Kit according to manufacturer’s instructions.

### Gene expression and RNAseq

Total RNA was isolated using the RNeasy Minikit (QIAGEN #74106; Hilden, Germany) and precipitated by the ethanol/sodium acetate method. RNA concentration and quality were initially measured using a Nanodrop 2000 (Thermofisher). Secondary analysis of concentration and quality was conducted by the Genomics CoRE Facility at the Icahn School of Medicine at Mount Sinai using Qubit RNA BR Assay Kit (ThermoFisher #Q10211). Library construction and RNA sequencing were performed by Novogene (Durham, NC). RNAseq data were processed using the NGS-Data-Charmer pipeline. Briefly, adaptors and low-quality bases were trimmed from reads, which were then aligned to the mm10 genome using HISAT2 (version 2.2.1). Read counts in the mm10 GENCODE annotation (version M22) were generated using FeatureCounts DESeq2 (version 1.24.0, R version 3.6.1) was then used to calculate differential expression from the read counts. The R package ‘biomaRt’ (version 2.40.5) was used to translate ensembl ids into common gene symbols.

### Gene ontology

Mouse genes involved in inflammatory processes were extracted from the Mouse Genome Informatics group database (MGI). MGI mouse genes with the GO term “Inflammatory responseL were extracted. Ingenuity Pathway Analysis was applied to all differential genes identified in the previous analysis.

### Quantitative reverse transcription PCR

For qPCR analysis, gene expression was measured in 4 replicates by Power UP SYBR Green Master Mix (ThermoFisher #A25778) using an ABI PRISM 7900HT Sequence Detection System. Hypoxanthine phosphoribosyl transferase (*Hprt*) expression level was used as an internal control and data was normalized using the 2^-ΔΔCt^ method [18]. Levels of target gene mRNA were expressed relative to those of GFP + Veh mice for *in vivo* studies. Primers used in this study were designed using Primer-BLAST software [19] and are listed in Table S1.

### Tissue processing

Four mice from each group (GR_100_ and GFP) were processed for sequencing in cerebral cortex (CTX) per mouse. After assessing sample quality, one FC sample was excluded due to poor quality, and all tissue samples from a single mouse were removed due to sample misclassification.

### Statistical Analysis

All figure values are presented as the mean and standard error of the mean (s.e.m.). Statistical tests are indicated in the figure legends. A confidence interval of 95 % was used for all analyses. In all studies, outliers (> 2 SD from the mean) were excluded. All statistical analysis was performed using GraphPad Prism 9 software (GraphPad Software, San Diego CA). \**p* < 0.05, \*\**p* < 0.01, \*\*\**p*<0.001, \*\*\*\**p*<0.0001, *ns* not significant. Statistically insignificant trends are indicated by *p*-value.

## Results

### 1. GFP-(GR)_100_ animal model of c9FTD/ALS exhibits behavioral impairment, neurodegeneration, and neuroinflammation, leading to a significant innate immune-driven response

To investigate the effects of GR_100_ induced neuronal stress in the brain in c9FTD/ALS, neonatal intracerebroventricular injections of adeno-associated virus serotype 9 (AAV_9_) vectors containing either GFP (control) or 100 repeats of the GR dipeptide (GFP-(GR)_100_), was conducted, as previously reported [20]. AAV_9_ has a strong expression pattern in neurons of the brain, but not microglia [21], allowing for the targeted investigation of neuronal-induced effects. The body weight of GFP and GFP-(GR)_100_ mice did not differ throughout the study (**supplementary Fig 1a**). Examination of c9FTD/ALS behavioral features in GFP- (GR)_100_ revealed significant mortality over the course of 3 months after injection, with over 25 % of animals dying before completion (**Fig 1a**). To assess behavioral responses, mice were tested for contextual memory, motor function, and anxiety. GFP-(GR)_100_ animals exhibited a significant reduction in freezing in the contextual memory paradigms (**\*\**p*<0**.**01, Fig 1b**), as well as a significant increase in falls in the hanging wire test (**\*\*\*\**p*<0**.**0001, Fig 1c**). Similarly, GFP-(GR)_100_ mice exhibited a significant elevated anxiety in the open field test (**\* *p*<0**.**05, Fig 1d**).

**Figure 1.**
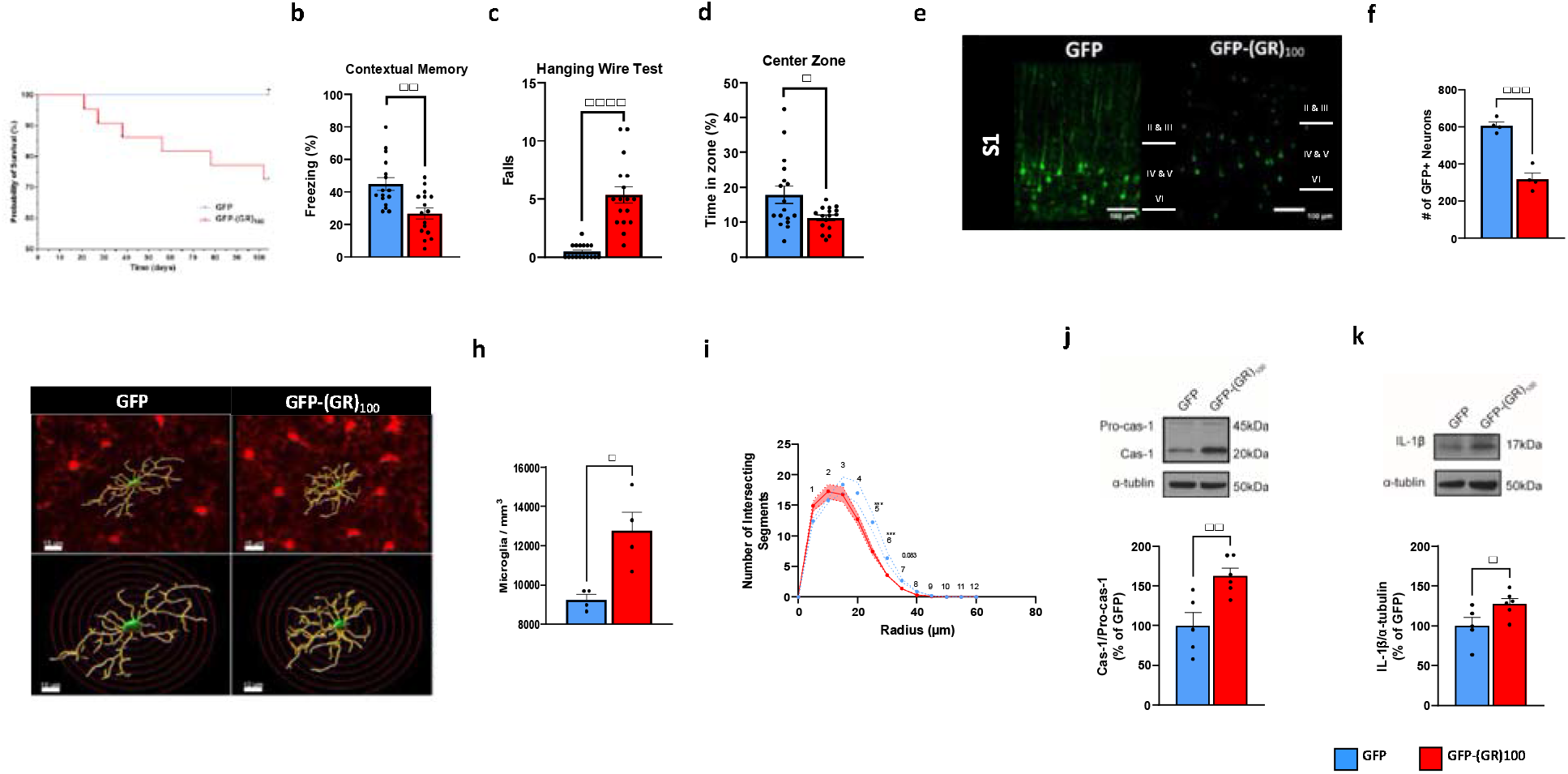
Innate immune-driven neuroinflammation and neurodegeneration in a GFP-(GR)_100_ model of c9FTD/ALS. (a) Probability of survival of GFP and GFP-(GR)_100_. (b) Freezing in contextual fear memory task. (c) Number of falls on the third day of the Hanging Wire Test. (d) Time spent in the central zone in the Open Field Test. (e) Representative GFP and GFP-(GR)_100-_expression neurons in the layers of IV and V of S1 cortical region. Green: GFP and GFP-(GR)_100_. Length bar, 100 µm. (f) Count of GFP- and GFP-(GR)_100_-expression neurons in the S1 cortical region. (g) 3D reconstruction of microglia in the S1 cortical region, with increasing radii in 5 µm steps to measure branch intersections. Length bars, 15 µm and 10 µm in top and bottom panels, respectively. Red: IBA-1 immunostaining. Green: Soma. Yellow: branches. (h) Microglial density in the S1 cortical region. (i) Sholl analysis of microglia in S1 cortical region. (j) Representative bands (upper panel) and densitometric quantification of the ratio of active caspase-1 to pro-caspase-1 (bottom panel) in the cerebral cortex, normalized to GFP. (k) Representative bands (upper panel) and densitometric quantification of cleaved IL-1β in the cerebral cortex (bottom panel), normalized to GFP. The probability of survival (a) was analyzed by log-rank test. For (b-d, f, h-k), * *p*<0.05, \*\**p*<0.01, \*\*\**p*<0.001, \*\*\*\**p*<0.0001 by unpaired *t*-test. Data are presented as mean ± SEM.

Next, cortical neuron degeneration was assessed by quantification of GFP-expressing neurons in the layers of IV and V in the S1 cortical region, which revealed a significant reduction in GFP-expressing neurons in GFP-(GR)_100_ mice compared to GFP mice (**\*\*\**p*<0**.**001, Fig 1f**). Microglia-mediated neuroinflammation has been described in cases of ALS [22]. Microgliosis has also been previously described to occur in this model, beginning at 1.5 months of age [23]. We hypothesized that microglia, the innate immune cells of the brain, may be causally related to the pathogenesis of c9FTD/ALS. To determine microglial response to the disease process, we first stereologically quantified the density of microglia in the S1, a relevant region of the brain that contributes to the behavioral impairments we observed in GFP-(GR)_100_. We found a significant increase of microglia density in GFP-(GR)_100_ mice compared to GFP mice (**\* *p*<0**.**05, Fig 1h**). Moreover, morphological analyses were conducted to assess activation state of microglia (**Fig 1g)**, which revealed a trending increase in the level of ramifications closer to the soma in GFP-(GR)_100_ mice as reflected by the increase in intersecting segments within 10 µm of the soma (**Fig 1g right top and bottom panels and Fig 1i)** compared to GFP mice (**Fig 1g left top and bottom panels and Fig 1i)**. This is accompanied by a statistically significant decrease in the number of intersecting segments further away from the soma (20 – 30 µm away from the soma) (**Fig 1i**) as assessed by Sholl analysis, indicative of an activated state with increased branch complexity as shown in **Fig 1g**. This increased density and morphological ramification of microglia is indicative of a neuroinflammatory response occurring in the S1 cortical regionally overlapping neuronal loss in GFP-(GR)_100_ mice.

Next, we investigated the effects of microglial activation seen in GFP-(GR)_100_ mice in neuroinflammatory responses as assessed by innate immunity including cleavage of caspase-1 and IL-1β level (**Fig 1j, 1k)**. Interestingly, we found that GFP-(GR)_100_ mice exhibited an increase in the ratio of active caspase-1 to pro-caspase-1 (**\*\**p*<0**.**01, Fig 1j**) in the cerebral cortex. Additionally, IL-1β was significantly increased in the same region of GFP-(GR)_100_ compared to GFP mice (**\* *p*<0**.**05, Fig 1k**). Collectively, the elevation of active caspase-1, pro-caspase-1, IL-1β, and activation of microglia suggests that there is a significant innate immune-driven response *in vivo* in GFP-(GR)_100_ mice potentially associated to activation of inflammasome in microglia.

### 2. Genetic ablation of *Nlrp3* inflammasome confers protection against innate immune-driven inflammation in the cerebral cortex of c9FTD/ALS mice

Based upon our results identifying significant innate immune-related inflammation in a GFP- (GR)_100_ model of c9FTD/ALS, we targeted the NLRP3 inflammasome complex via genetic ablation. To test the effects of genetic ablation of *Nlrp3*, GFP-(GR)_100_ or GFP expression in the mouse brain we conducted neonatal intracerebroventricular injections of AAV_9_ vectors in mice lacking the *Nlrp3* gene (*Nlrp3*^−/−^).

Weights of *Nlrp3*^−/−^-GFP and *Nlrp3*^−/−^-GFP-(GR)_100_ did not differ over the course of the study (**supplementary Fig 1b**). Next, examination of the behavioral effects of inflammasome targeted interventions revealed no change in mortality in *Nlrp3*^−/−^-GFP-(GR)_100_ micec ompared to *Nlrp3*^−/−^-GFP mice (**Fig 2a**). Assessment of memory function demonstrated unaffected contextual memory in *Nlrp3*^−/−^-GFP-(GR)_100_ mice compared to *Nlrp3*^−/−^-GFP mice (**Fig 2b**). Similarly, motor function (**Fig 2c**) and anxiety (**Fig 2d**) assessed by hanging wire and open field tests remained unaffected in *Nlrp3*^−/−^-GFP-(GR)_100_, with mice equally performing as found in *Nlrp3*^−/−^-GFP mice.

**Figure 2.**
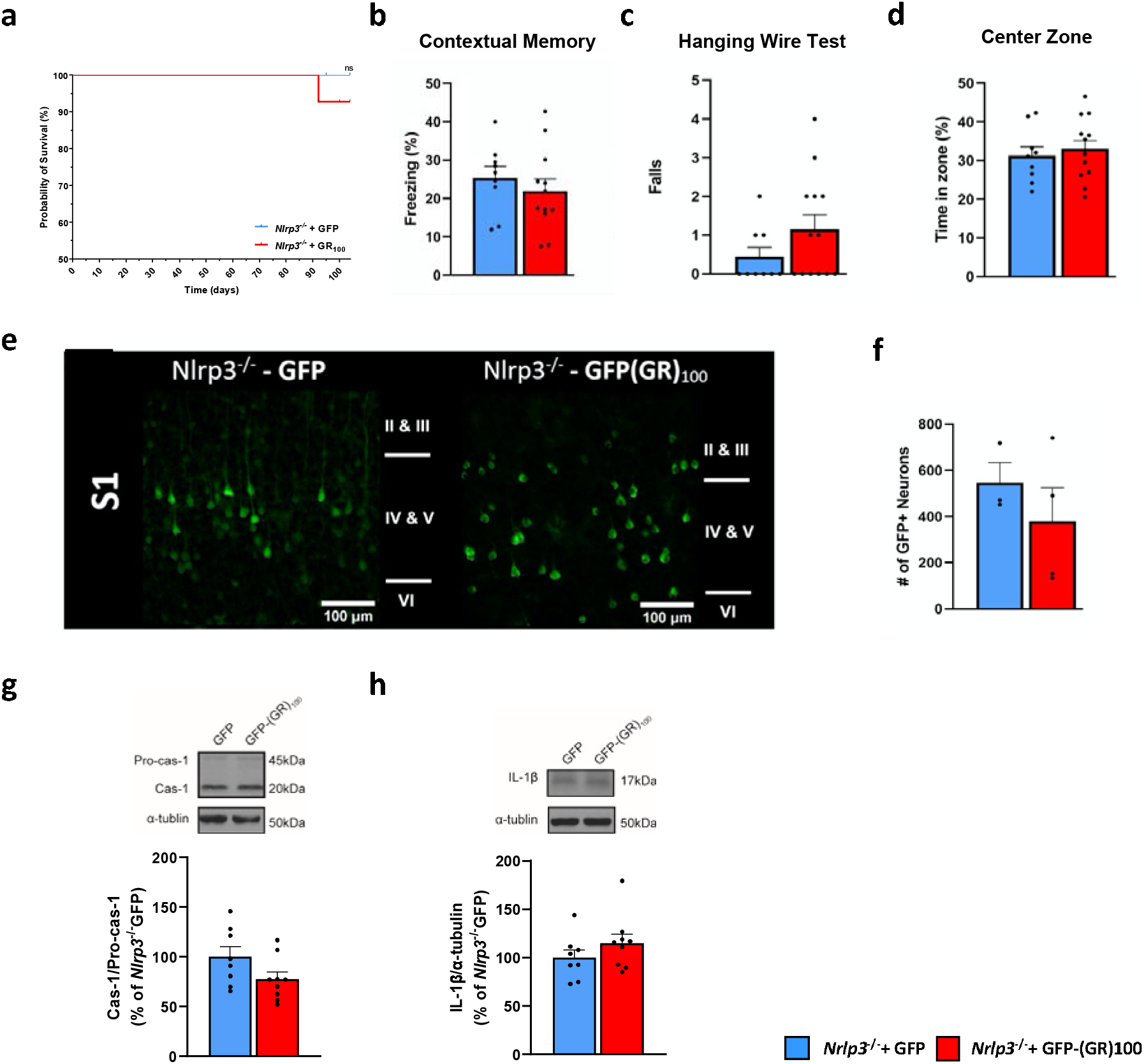
Genetic ablation of NLRP3 inflammasome attenuates neurodegeneration and behavioral deficits in a GFP-(GR)_100_ model of c9FTD/ALS. (a) Probability of survival of *Nlrp3*^−/−^-GFP and *Nlrp3*^−/−^-GFP(GR)_100_mice. (b) Freezing in contextual fear memory task. (c) Number of falls in the third day of the Hanging Wire Test. (d) Time spent in the central zone in the Open Field Test. **(e)** Representative *Nlrp3*^−/−^-GFP and *Nlrp3*^−/−^-GFP(GR)_100_ expressing neurons in the layers IV and V of S1 cortical region, Green: GFP and GFP-(GR)_100_. Length bar, 100 µm. (f) Count of *Nlrp3*^−/−^-GFP- and *Nlrp3*^−/−^-GFP-(GR)_100_-expression neurons in the layers IV and V of S1 cortical region. (g) Representative bands (upper panel) and densitometric quantification of the ratio of active caspase-1 to pro-caspase-1 (bottom panel) in the cerebral cortex, normalized to GFP. (h) Representative bands (upper panel) and densitometric quantification of the ratio of cleaved IL-1β caspase-1 to Tubulin (bottom panel) in the cerebral cortex, normalized to *Nlrp3*^−/−^-GFP. The probability of survival (a), was analyzed by log-rank test. For (b-d, f, g, h), were analyzed by unpaired *t*-test. Data are presented as mean ± SEM.

We next investigated the effects of targeting the NLRP3 inflammasome on neurodegeneration in GFP-(GR)100 mice in the S1 cortical region. No changes in the count of GFP-expressing neurons were noted in *Nlrp3*^−/−^-GFP-(GR)_100_ compared to *Nlrp3*^−/−^-GFP (**Fig 2e, f**). Lack of neurodegeneration and protection in the associated behavioral tasks indicate a role of *Nlrp3* in c9FTD/ALS pathogenesis.

Based our evidence inflammasome-mediated neuroinflammation in the cereberal cortex of WT GFP-(GR)_100_ mice (**Fig 1j and 1k**), we further investigated these findings in the cereberal cortex of *Nlrp3*^−/−^-GFP-(GR)_100_ mice. We found that no change in the ratio active caspase-1 to pro-caspase-1 in *Nlrp3*^−/−^-GFP-(GR)_100_ mice compared to *Nlrp3*^−/−^-GFP mice (**Fig 2g**). Similarly, no significant differences in level of active IL-1β were observed in *Nlrp3*^−/−^-GFP- (GR)_100_ mice compared to *Nlrp3*^−/−^-GFP mice (**Fig 2h**). This evidence supports hypothesis of microglia-driven innate immune responses in the brain of c9FTD/ALS.

### 3. GFP-(GR)_100_ expression induces transcriptional changes in innate immune microglia-mediated responses of c9FTD/ALS animals

We next investigated the transcriptional regulation of genes in the CTX of GFP-(GR)_100_ mice relative to GFP mice to assess regional distribution of changes. Overall transcriptional patterns depicting the significantly differentially expressed genes are visualized in **Fig 3a**, with the top three mapped upregulated and downregulated genes labeled as identified by the magnitude of Log_2_Fold Change, as visualized in a volcano plot. The top-upregulated mapped gene in GFP-(GR)_100_ is *Cxcl10* (Log_2_fold Change = 4.51; *p* _adj_ = 0.06, **Fig 3a**), which has been previously shown to result in the proliferation and activation of microglia [24]. Based on this *in vivo* data suggesting a causal link between inflammasome activation and neurodegeneration resulting in behavioral impairments, we hypothesized that GFP-(GR)_100_ expression would invoke transcriptional changes in innate immune responses. We proceeded with the examination of the pathways and regulators enriched in the differentially expressed genes using Ingenuity Pathway Analysis (IPA) confirms enrichment in neuroinflammatory signaling pathways including ‘*Complement System’, ‘TREM1 Signaling’, ‘Chemokine Signaling’, and ‘NF-*κ*B Signaling’*, in the CTX (**Fig 3b**). These pathways are core regulators of inflammatory responses and microglial activation. Amongst validated genes (**Fig 3c**), we found that *Cxcl10* is significantly upregulated, with a 7-fold increase in GFP-(GR)_100_ mice, compared to GFP mice (**Fig 3d**). The transcriptomic changes exhibited in GFP-(GR)_100_ animals heavily point to significant innate immune microglia-mediated responses. Our findings of significant *Cxcl10* upregulation suggest there is a neuronal signal released that can activate microglia and induce innate immune cascades in GFP-(GR)_100_ mice.

**Figure 3.**
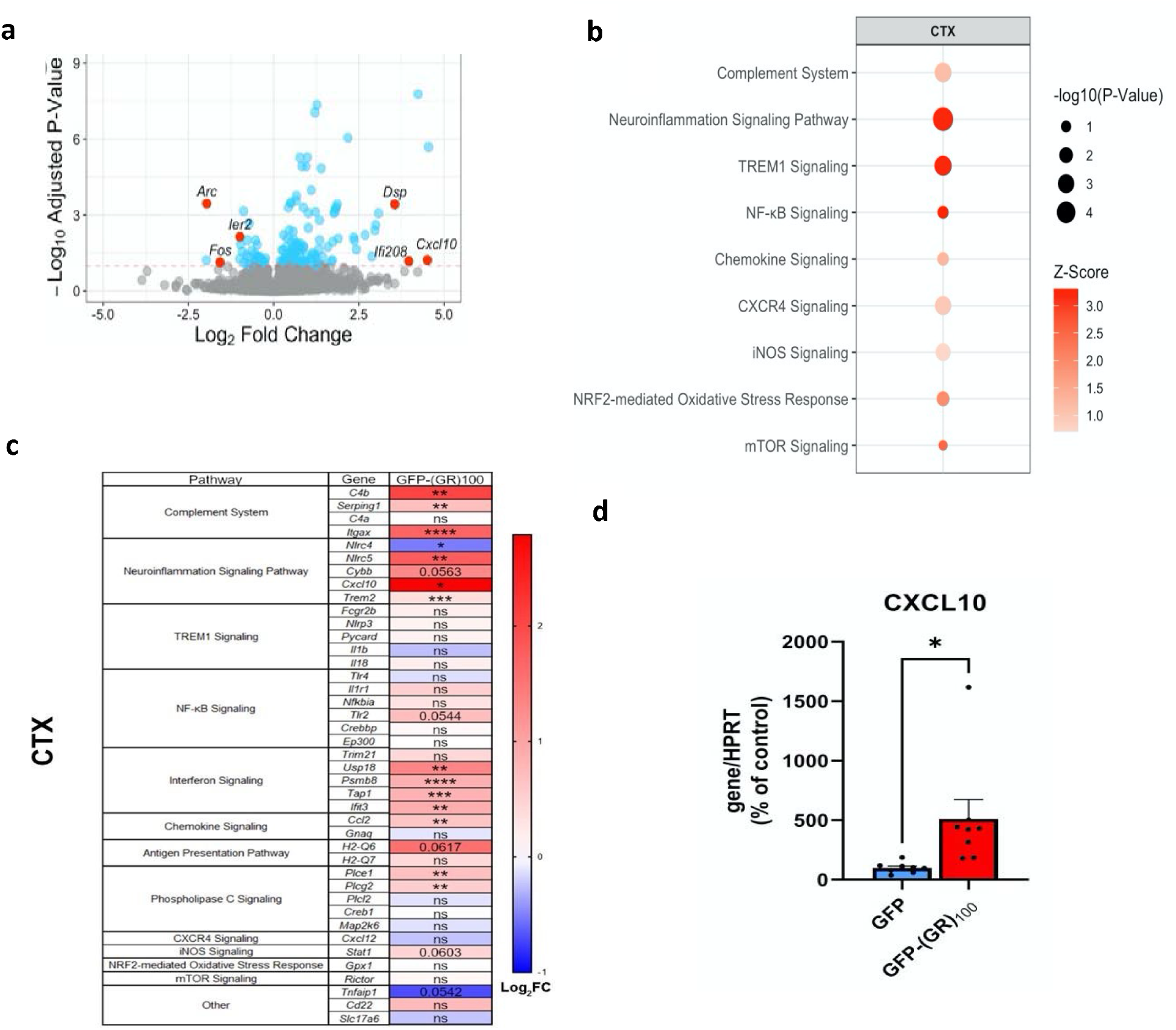
Transcriptional changes of GFP-(GR)_100_. (a) A volcano plot of CTX transcription. The X-axis represents the log2 fold change, while the Y-axis represents the negative log10 FDR-adjusted *p*-value. Genes with an adjusted *p*-value less than 0.10 are shown in blue. The 3 most highly upregulated mapped genes and 3 most highly downregulated genes are highlighted in red. (b) Inflammation-related IPA pathway; select immune system-related pathways are shown for the CTX. The *p*-value of each entry is represented by the size of the corresponding circle. The z-score, indicating the computed activation or repression of the given pathway, is indicated with a color gradient. (c) Validation of significant IPA genes identified via RNAseq. Gene ontologies are indicated on the left. Colors indicate log_2_ fold change (Log_2_FC). In c, \**p*<0.05, \*\**p*<0.01, \*\*\**p*<0.001, \*\*\*\**p*<0.0001 by Two-Tailed Welch’s t-test. Statistically insignificant trends (*p*<0.1) are indicated. *ns*: not significant (*p* > 0.1). (d) Validation of the most highly upregulated gene, *Cxcl10* mRNA expression, in the CTX of GFP and GFP-(GR)_100_ mice. In d, * *p*<0.05 by Two-tailed Welch’s t Test.

### 4. GFP-(GR)_100_ induces secretion of neuronal CXCL10 activating a microglial innate immune response

As *Cxcl10* is an important signal for the immune responses of microglia [24], we investigated the effect of GFP-(GR)_100_ infection in primary neuronal and microglial cultures on CXCL10 production, and whether CXCL10 induces microglia activation. First, we investigated whether the expression of GFP-(GR)_100_ induced neuronal cell death *in vitro*. GFP-(GR)_100_ transfected neurons exhibited a significant increase in the release of lactate dehydrogenase (LDH) compared to GFP transfected neurons (**\**p*<0**.**05, Fig 4a**), confirming the GFP-(GR)_100_-induced neuronal injury and death observed *in vivo*. As *Cxcl10* was shown to be highly upregulated *in vivo* in GFP-(GR)_100_ mice, we hypothesized that neurons are able to produce and secrete CXCL10, in response to GFP-(GR)_100_. To test this, primary cortical neurons were transfected with either AAV_9_-GFP or AAV_9_-GR_100_ and cell supernatants were analyzed for CXCL10 concentration. GFP-(GR)_100_ neurons demonstrated significantly increased production of CXCL10 compared to GFP transfected neurons (**\*\*\*\**p*<0**.**0001, Fig 4b**). As we identified *Cxcl10* is highly upregulated *in vivo* and was determined experimentally to be significantly released by primary cortical GFP-(GR)_100_ neurons *in vitro* (**Fig 4b**), we next investigated whether CXCL10 activates microglia.

**Figure 4.**
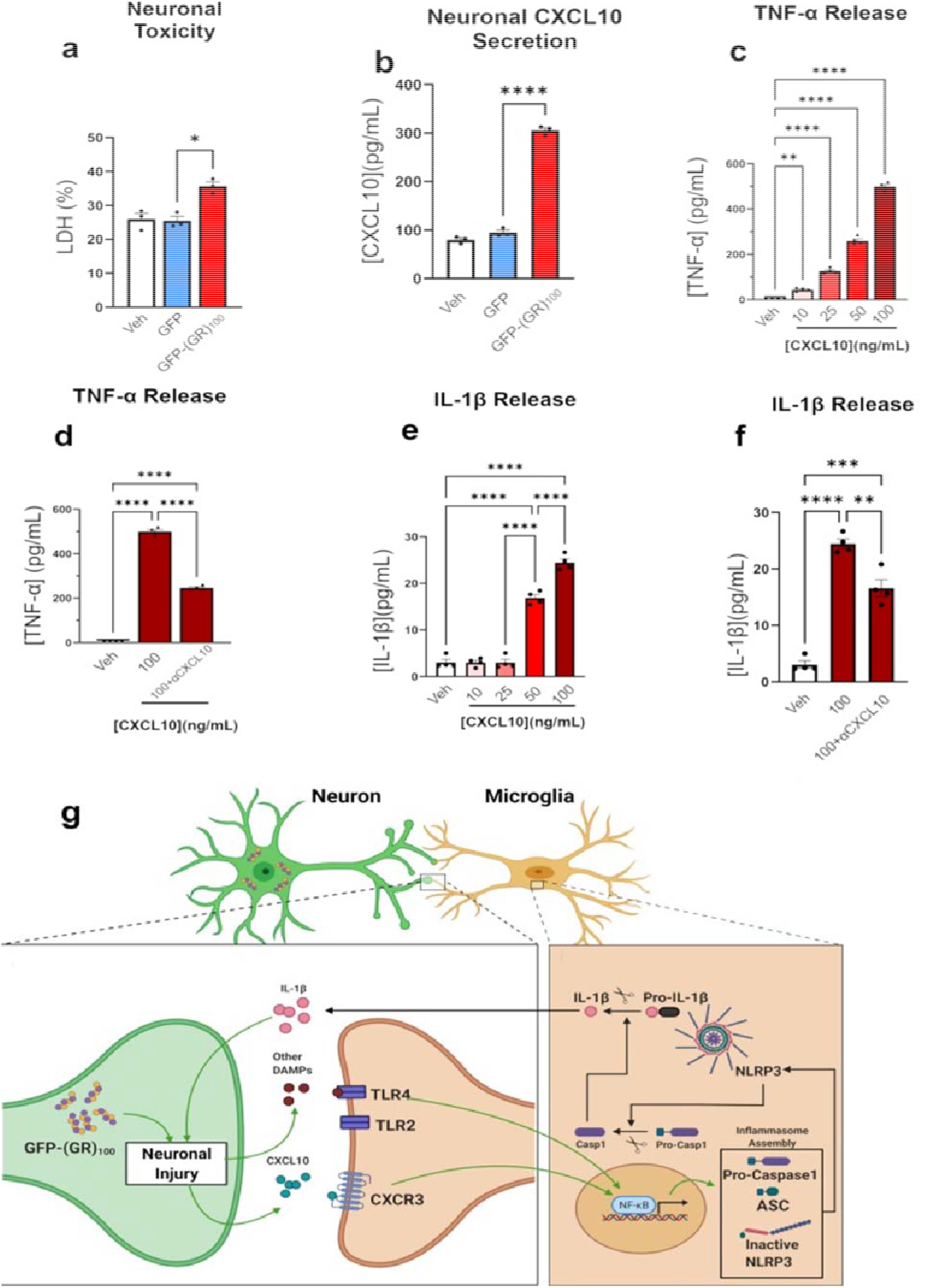
In vitro evaluation of neuronal signaling and resultant activation microglia in c9FTD/ALS. (a) Evaluation of neurotoxicity of GFP-(GR)_100_. (b) Neuronal release of CXCL10 after transfection with GFP-(GR)_100_. (c) CXCL10 mediated dose-dependent release of TNF-α. (d) Neutralization of CXCL10 and TNF-α release. (e) CXCL10 mediated dose-dependent release of IL-1β. (f) Neutralization of CXCL10 and IL-1β release. (g) Schematized mechanism of GFP-(GR)_100_ inducing the neuronal release of CXCL10, which may then bind to microglial receptors and initiate an inflammasome response. The data shown correspond to average values of biological triplicates ran in technical duplicates. For a-f, \**p*<0.05, \*\**p*<0.01, \*\*\**p*<0.001, \*\*\*\**p*<0.0001 by one-way ANOVA.

Tumor necrosis factor-alpha (TNF-α) constitutes a hallmark for microglia activation and has previously been shown to play an important role in the transcriptional regulation of components of the NLRP3 inflammasome [25]. To test the ability of CXCL10 to induce TNF-α expression, primary cortical microglia cultures were treated with increasing CXCL10 concentration and assessed released level of TNF-α in cell supernatant. We found that CXCL10 significantly increased the microglial production of TNF-α in a dose-dependent manner (**\*\**p*<0**.**01; \*\*\*\**p*<0**.**0001, Fig 4c**), while the co-treatment with a CXCL10 neutralizing antibody results in a significant reduction in the TNF-α secreted by microglia (**\*\*\*\**p*<0**.**0001, Fig 4d**). Finally, to analyze if CXCL10 mediates the activation of IL-1β processing inflammasome complexes, we measured the release of IL-1β from primary cortical microglia in response to increasing concentration of CXCL10. We found a significant, dose-dependent increased secretion of IL-1β (**\*\*\*\**p*<0**.**0001, Fig 4e**). Co-treatment with CXCL10 neutralizing antibody results in a significant reduction in IL-1β released by microglia (**\**p*<0**.**05; \*\**p*<0**.**01; \*\*\*\**p*<0**.**0001, Fig 4f**). This evidence suggests neuronal-microglial crosstalk whereby GFP-(GR)_100_ induces secretion of neuronal CXCL10, activating a microglial innate immune response, resulting in inflammasome activation and subsequent neuroinflammation.

In conjunction with the rest of our data, these experiments suggest that neurons in c9FTD/ALS produce damage associated molecular patters (DAMPs) such as CXCL10 caused by GFP-(GR)_100_, which significantly stimulates pro-inflammatory reactions from microglia, leading to inflammasome activation (**Fig 4g**).

## Discussion

Our study provides evidence linking inflammasome activation to neurodegeneration and provides a basis for the investigation of innate immune inflammasome inhibitors as a treatment of disease in c9FTD/ALS. Here, we identify a key mechanism by which neuronal stress initiated by G_4_C_2_ HREs in c9FTD/ALS can activate microglia, producing an inflammasome-dependent innate immune response, which can be therapeutically targeted. As neuronal stress occurs throughout various forms of FTD/ALS [26–28], the self-perpetuating cycle of neuronal stress inducing microglial inflammasome activation and resultant neuroinflammation may represent a conserved therapeutic substrate throughout the various disease subtypes of FTD/ALS. Ultimately, we demonstrate that by targeting microglial reactivity through inflammasome inhibition, FTD/ALS pathogenesis was significantly attenuated through the genetic ablation of *Nlrp3*.

The results of these studies identify the contributions of inflammasome activation in microglia to the pathogenesis of c9FTD/ALS. Microglial activation and the activation of the NLRP3 inflammasome have been demonstrated as crucial mediators in other neurodegenerative conditions including Alzheimer’s disease [20], Parkinson’s disease [29], and primary progressive multiple sclerosis [30]. Inflammation has also been noted pre-symptomatically in models of ALS [31,32] and the incidence of ALS has been demonstrated to be higher in individuals who have been diagnosed with autoimmune disease [33], indicating a putative role in the disease pathogenesis and a target for therapeutic intervention. Neuroimaging studies conducted on individuals with ALS have demonstrated microglial activation throughout the brain [22,34].

Upon examination of the microglia in the S1 cortical region, we observed microgliosis consistent with previously identified findings [23]. Morphologically, microglia in GFP-(GR)_100_ were in an activated state; these findings are congruent with those identified in other neurodegenerative diseases, including Alzheimer’s disease, in which morphologically activated microglia are present [35]. Inflammasome activation has been identified in other forms of ALS [36–38]. We noted an increase in the ratio of active-caspase-1 to pro-caspase-1 in the CTX of WT GFP-(GR)_100_ mice; these increases were attenuated via genetic ablation of *Nlrp3* as well as the production of active IL-1β. These results shed light on the role the NLRP3 inflammasome might have on the pathogenesis of FTD/ALS.

In addition to improvements in neurodegeneration and neuroinflammation, full genetic ablation of the *Nlrp3*^−/−^ inflammasome conferred significant protection behaviorally in *Nlrp3*^−/−^-GFP-(GR)_100_ mice compared to *Nlrp3*^−/−^-GFP mice; mortality was almost completely eliminated, contextual memory was preserved, motor function was protected, and no increase in anxiety was observed. To determine if the NLRP3 inflammasome can be therapeutically targeted, further studies may employ experimental models whereby *Nlrp3* ablation by Cre dependent-designer receptor exclusively activated by designer drugs (DREADD) mice allow for temporal control of *Nlrp3* expression, thereby evaluating both its role in pre-symptomatic disease and evaluating whether the NLRP3 inflammasome may serve as a therapeutic target after symptom onset.

*Cxcl10*, which we found to be highly upregulated in the cerebral cortex of GFP-(GR)_100_ mice, encodes for a small cytokine belonging to the CXC chemokines family. Upon neuronal death, CXCL10 is produced and exerts a chemotactic function by attracting microglia and CD8^+^ T cells, and can induce the activation of microglia, thereby causing the release of pro-inflammatory cytokines [39]. Chemotactic effects are demonstrated in our FTD/ALS model by the microgliosis in the cortex of GFP-(GR)_100_ mice where we see a significant increase in the density of microglia. Neuronal stress has been demonstrated in other forms of ALS, including in cases of sporadic ALS [40]. Dysregulation of CXCL10 chemotaxis in peripheral blood cells from ALS patients has been directly observed, which was associated with increased inflammatory responses [41]. This evidence supports our *in vitro* findings, where we demonstrated that neuronal cultures expressing (GR)_100_ dipeptides released higher concentrations of CXCL10, and the treatment of microglia cultures with increasing concentrations of CXCL10 promoted a dose-dependent increase in TNF-α and in IL-1β production indicating microglial activation.

In summary, our studies identify a novel neuronal-microglial crosstalk mechanism in c9FTD/ALS whereby neuronal stress-induced secretion of CXCL10 triggers inflammasome activation in microglia, thereby creating a self-propagating cycle of neurodegeneration and neuroinflammation. Our findings show targeting the inflammasome responses represents a putative therapeutic strategy, as evidenced by the genetic ablation of *Nlrp3* and resultant protection from behavioral impairment and neuropathologies.

## Supplementary Information

**Supplementary Table S1.**
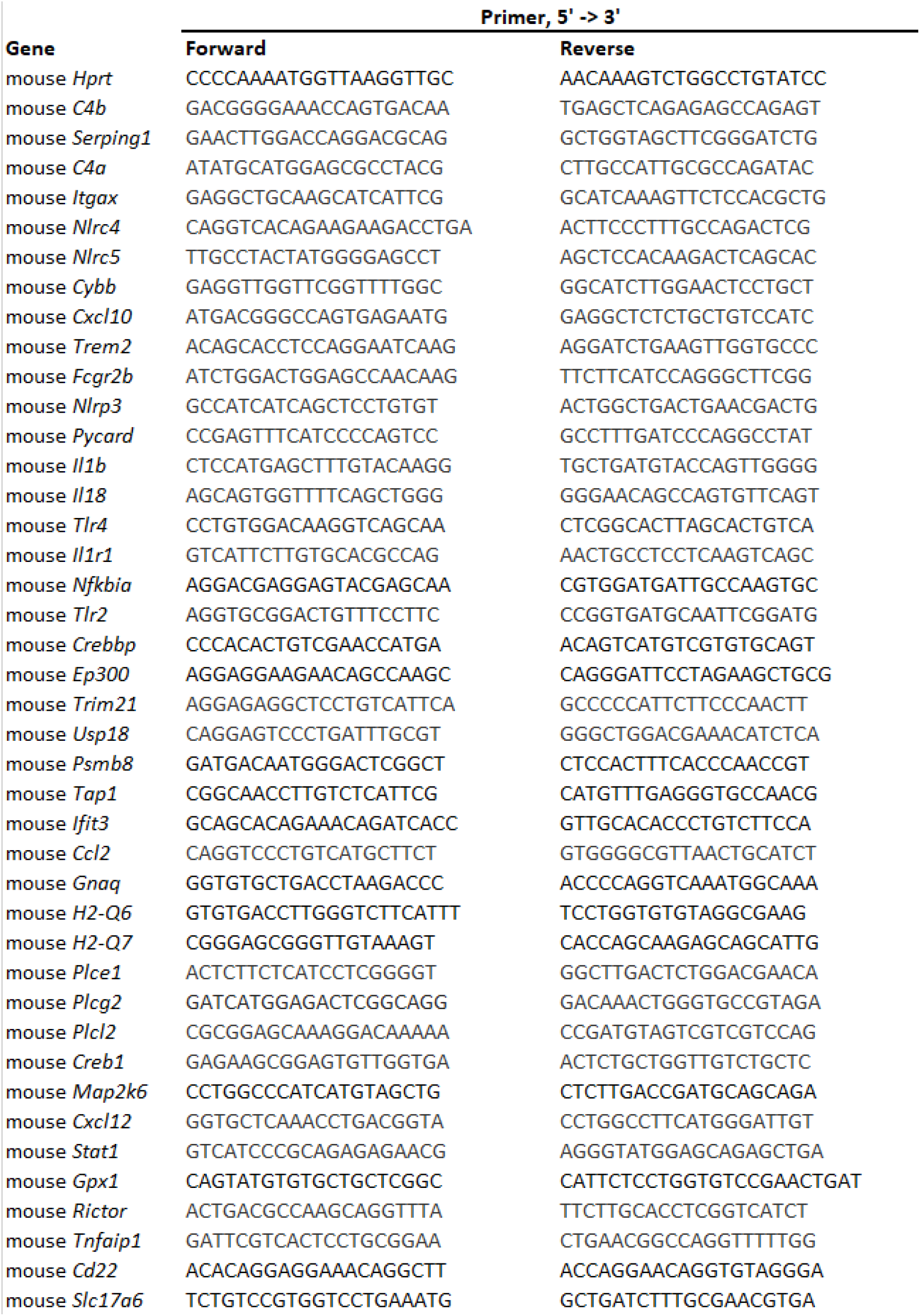
Primers used in this study.

**Supplementary Figure 1.**
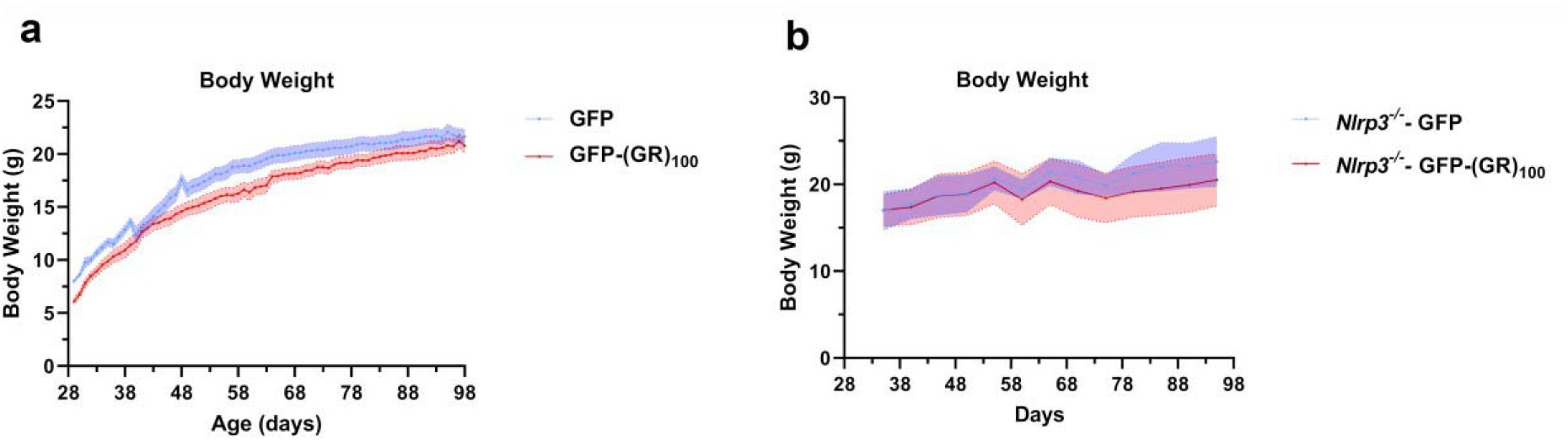
Physiological monitoring data of GFP and GFP-(GR)_100_ (a) and *Nlrp3*^−/−^-GFP and *Nlrp3*^−/−^-GFP(GR)_100_ mice (b).

## Ethics approval

All of the experimental procedures were approved by the Mount Sinai Institutional Animal Care and Use Committee (IACUC) (Approval number: IACUC-2019-0006).

## Consent to participate

Not applicable.

## Consent for publication

Not applicable

## Availability of data and materials

The authors declare that the data supporting the findings of this study are available within the paper and its Supplementary information files. Any remaining data that support the results of the study will be available from the corresponding author upon reasonable request.

## Competing interests

The authors declare that they have no conflicts of interest with the contents of this article.

## Funding

The study was supported by the generous support of the Altschul Foundation to G.M.P. and by the Veteran Administration program on ALS. G.M.P. holds a Senior VA Career Scientist Award. We acknowledge that the contents of this study do not represent the views of the NCCIH, the ODS, the National Institutes of Health, the U.S. Department of Veterans Affairs, or the United States Government.

## Author Contributions

K.J.T and C.S conducted *in vivo* experiments of the GFP-(GR)_100_ model. Immunofluorescence staining, image acquisition, and analysis were performed by K.J.T, C.S, U.H.I, U.R, and T.O. *In vitro* experiments were conducted by M.S.V, M.A.R, and R.I.A. Molecular analyses, including Western Blots, and RT-qPCR were performed by K.J.T, C.S, E.Y, M.S.V., H.W, and M.A.R. Bioinformatics analyses were conducted by H.W and M.E. Generation and purification of the GFP-(GR)_100_ and GFP viruses were completed by Y.J.Z and L.P. K.J.T, C.S, F.J.H, and G.M.P designed the project. K.J.T, C.S, R.I.A, E.Y, and G.M.P prepared the manuscript. All authors discussed and commented on the article.

## Acknowledgments

We thank Josh Palmieri for his impeccable work with administrative duties.

## Notes

### Competing Interest Statement

The authors have declared no competing interest.

### Summary of Updates

We revised all figures and associated with description in the manuscript.

